# A time-course study of actively stained mouse brains: DTI parameter and connectomic stability over one year

**DOI:** 10.1101/2020.12.02.407338

**Authors:** Jaclyn Xiao, Kathryn J. Hornburg, Gary Cofer, James J. Cook, Yi Qi, G Allan Johnson

## Abstract

While the application of diffusion tensor imaging (DTI), tractography, and connectomics to fixed ex-vivo tissue is a common practice today, there have been limited studies examining the effects of fixation on brain microstructure over extended periods. This time-course study reports the changes of regional brain volumes and diffusion scalar parameters, such as fractional anisotropy across twelve representative brain regions as measures of brain structural stability. The scalar DTI parameters and regional volumes were highly variable over the first two weeks after fixation. The same parameters were stable over a two to eight-week window after fixation which means confounds from tissue stability over that scanning window are minimal. Quantitative connectomes were analyzed over the same time period with extension out to one year. While there is some change in the scalar metrics at one year after fixation, these changes are sufficiently small, particularly in white matter to support reproducible connectomes over a period ranging from two weeks to one year post fixation. These findings delineate a stable scanning period during which brain volumes, diffusion scalar metrics and connectomes are remarkably stable.

## INTRODUCTION

The use of diffusion tensor imaging (DTI) has grown over the past few years, spawning the development of technical advancements^1^ and numerous applications in connectomics^2–4^. In multiple human populations studies, eg. the human connectome project, the ENGIMA consortium and the ADNI initiative, DTI has proven particularly useful in understanding human cognition, teasing out genetic differences and understanding pathology ^5–8^. The application of DTI has expanded into animal studies, where the scalar parameters have proven useful in understanding of structural changes of the brain during maturation changes that are genetically driven and changes due to disease or injury ^9 10^.

The development of disease models and the genetic control that mouse studies offer provide a controlled approach to the understanding of disease mechanisms not possible in a clinical setting. The postmortem mouse allows for a level of imaging precision otherwise unachievable in human or live animal studies, since one can employ specialized staining methods and long scan times^11–13^. However, the study of the postmortem tissue brings its own unique challenges. Diffusion in unfixed postmortem tissues change over time due to autolysis^14^. Fixation, which affects water diffusivity in tissue and promotes cross-linking of proteins^15^, can be used to preserve and stabilize tissue architecture. Consequently, the errors associated with a postmortem study, like tissue decomposition, can be minimized.

Active staining introduces fixative and contrast agent through direct cardiac infusion of the live animal minimizing tissue destruction from autolysis^12,16,17^. Active staining reduces the spin lattice relaxation time (T1) through the use of a gadolinium contrast agent, leading to improved signal to noise, spatial resolution, and image contrast. However, until the fixative diffuses uniformly throughout the tissue, the brain remains susceptible to autolysis^15,18^. Crosslinking induced by fixation takes place on a time scale of minutes to years during which the relaxation and diffusion properties can undergo profound changes^18,19^. Furthermore, the effects of fixation on spin-spin relaxation (T2) over time post fixation^19–22^ can lead to additional uncertainties in DTI parameters in fixed tissue.

Previous studies have examined diffusion parameters in live versus ex vivo fixed tissue. Most have concluded that while the absolute values may change, relative values of MR parameters in fixed tissue (eg. ratio fractional anisotropy of gray matter to white matter) are similar to those in vivo^23–26^. However, few studies have looked at the variation of DTI parameters at extended time post fixation. Most reports suggest scalar DTI parameters (fractional anisotropy, FA; axial diffusivity, AD; and radial diffusivity, RD) remain unchanged over time post fixation^27–30^. The impact of post mortem interval (PMI), i.e. the time between death and fixation during which the tissue undergoes autolysis has been studied ^14^. But scan interval (SI), the time post fixation until scanning, has been seen as something of minimal impact^31^. Other studies have investigated the effects of SI on T1 and T2. Most of these studies found that the T2 decreases sharply within the immediate period following fixation – a period found to last a week - before reaching a plateau up to 1-year postmortem^20–22,30,32^. It could thus be hypothesized that the DTI parameters, which are affected by T1 and T2, should stabilize following an initial period after fixation, once the brain has fully fixed. As emphasized through these dichotomous hypotheses, the previous literature observations with regards to DTI stability over time have been mixed.

While studies have separately addressed the changes in tissue variables such as volume or diffusion metrics over time post fixation, no study has yet to cohesively analyzed all these metrics across a large and consistent set of time points to allow group comparative analysis. Furthermore, no study has investigated the effects of active staining perfusion fixation on parameters that contribute to connectome tracking nor analyzed statistical differences in connectomes across time.

Our current work flow for quantitative connectomics involves active staining of a batch (n=6-24) of animals at a time. Our scan protocol acquires 51 three-dimensional (3D) volumes requiring a total scan time per specimen of ~12 hrs. The practical consequence is that there exists a spread in the interval between perfusion and scanning. The goal of this study was to understand the sources of variability in actively stained mouse brain over time to establish a scan interval during which changes from fixation would have little effect on group comparisons of scalar DTI measures, volumes and connectomes.

## EXPERIMENTAL

All animal experimentation was performed in accordance with the National Institutes of Health Guidelines for Animal Care and Use of Laboratory Animals and the protocols of the Duke University Animal Care and Use Committee.

### Specimen Preparation

Eight male C57BL/6J adult (90 day) mice were acquired (The Jackson Laboratory, Bar Harbor, ME, Cat# 000664) for MR imaging. The mice were prepared using an active staining technique ^17^ in which the mice were transcardially perfused through the left heart ventricle with a solution of 10% ProHance (Bracco Diagnostics, Princeton NJ), and 10% buffered formalin. Following fixation, the heads were removed and placed in 10% buffered formalin for 24 hours to allow fixation to continue. At the end of 24 hours, the brains, still in the cranial vault were removed from the formalin, rinsed and immersed in a 0.5% ProHance/phosphate buffered saline (PBS) solution to restore the T2.

### Data Acquisition

All data were acquired on a 9.4-Tesla Agilent Direct Drive MRI system utilizing a Stesjkal-Tanner spin echo sequence^33^. Acquisition time was reduced using a compressed sensing algorithm with a compression factor of 8 ^34^. A 420 × 256 × 256 image matrix was acquired over an 18.9 × 11.52 × 11.52 mm^3^ field of view with a native isotropic spatial resolution of 45 μm^3^ using MRI scan parameters (repetition time (TR) = 100 ms; echo time (TE) = 12.7 ms, b-value = 4000 s/mm^2^). Forty-six diffusion-weighted 3D volumes were acquired with the gradient direction of each of volume uniformly distributed on the unit sphere. Five base line (b0) volumes were acquired. A post processing pipeline registered the five b0 volumes together to generate the average baseline. The diffusion weighted volumes were registered to this baseline to correct for eddy currents^13^.

MR images were gathered at seven time points post-fixation (2-3 days, 1 week, 2 weeks, 3 weeks, 4 weeks, 8 weeks, and nominally 1 year). The scan intervals for each specimen are shown in Figure 1. Two specimens were scanned for the 2-3 day, 2-week, 3-week, 4-week, and 8-week time period. Four specimens were scanned at all time points. Two additional specimens were scanned only at the 1-year time point.

**Figure 1:**
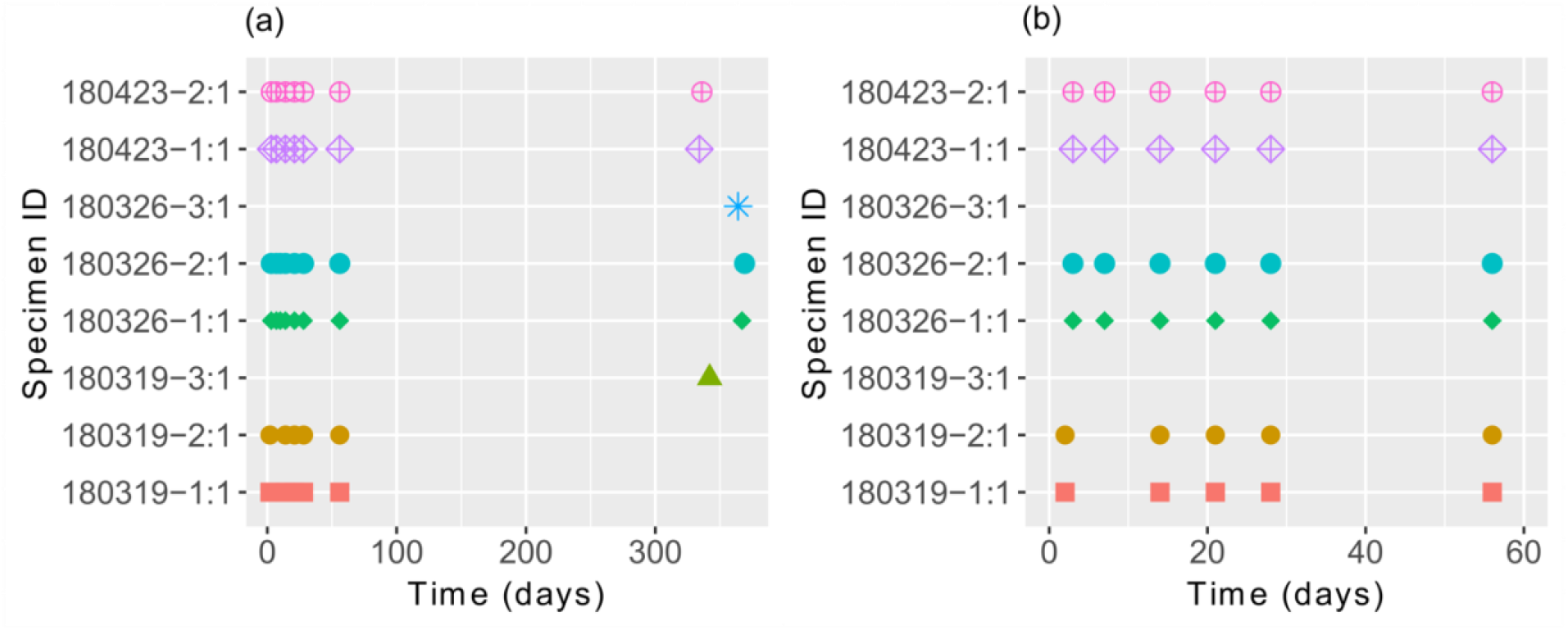
Scan time after perfusion for the 8 specimens used in this study: (a) all the time points (2-3 days to 1 year post fixation), and (b) detail of the time region from 2-3 days to 8 weeks post fixation.

### Atlas Mapping to MR Images

The SAMBA pipeline ^35^ was used to register the Waxhom Space (WHS) atlas^36^ with labels onto each specimen. The pipeline uses two steps to create the label mapping: an affine step using the diffusion weighted image (DWI) and diffeomorphic registration utilizing FA image. The WHS atlas includes 166 (symmetric) regions on each hemisphere in an adult C57BL/6J mouse which are mapped onto all the diffusion data in this study^36^. This provides ready extraction of the mean and standard deviation of the volume and diffusion scalar metrics from all 332 regions of interest in all the scans. It also provides the nodes used in defining the connectome for each specimen.

### Selection of Characteristic Regions

Twelve regions of interest (ROI) were followed across the entire time span; nine white matter and three gray matter regions. These regions are shown in Figure 2. The white matter regions: the anterior commissure (AC), corpus callosum (CC), cingulum (CG), fimbria (FI), internal capsule (IC), raphe nucleus (RN), cerebral peduncle (CP), middle cerebellar peduncle (MCP), and inferior cerebellar peduncle (ICP), were selected because of their relatively large volumes, to minimize partial volume errors, while still being representative for the whole brain. Some of these regions represent challenges with mapping: proximity to the ventricles (FI) and smaller volume (cerebellar peduncles). The grey matter regions: somatosensory cortex (S1), cingulate cortex (C), and primary visual cortex (V), were selected because these represent bulk cortical regions which also represent a challenge with automated mapping^37,38^. The contrast between cortical regions (eg. motor and sensory) is not particularly high in the MR images. In addition, layers within some cortical sections are not well delineated. So, we have combined some of these regions to simplify the segmentation.

**Figure 2:**
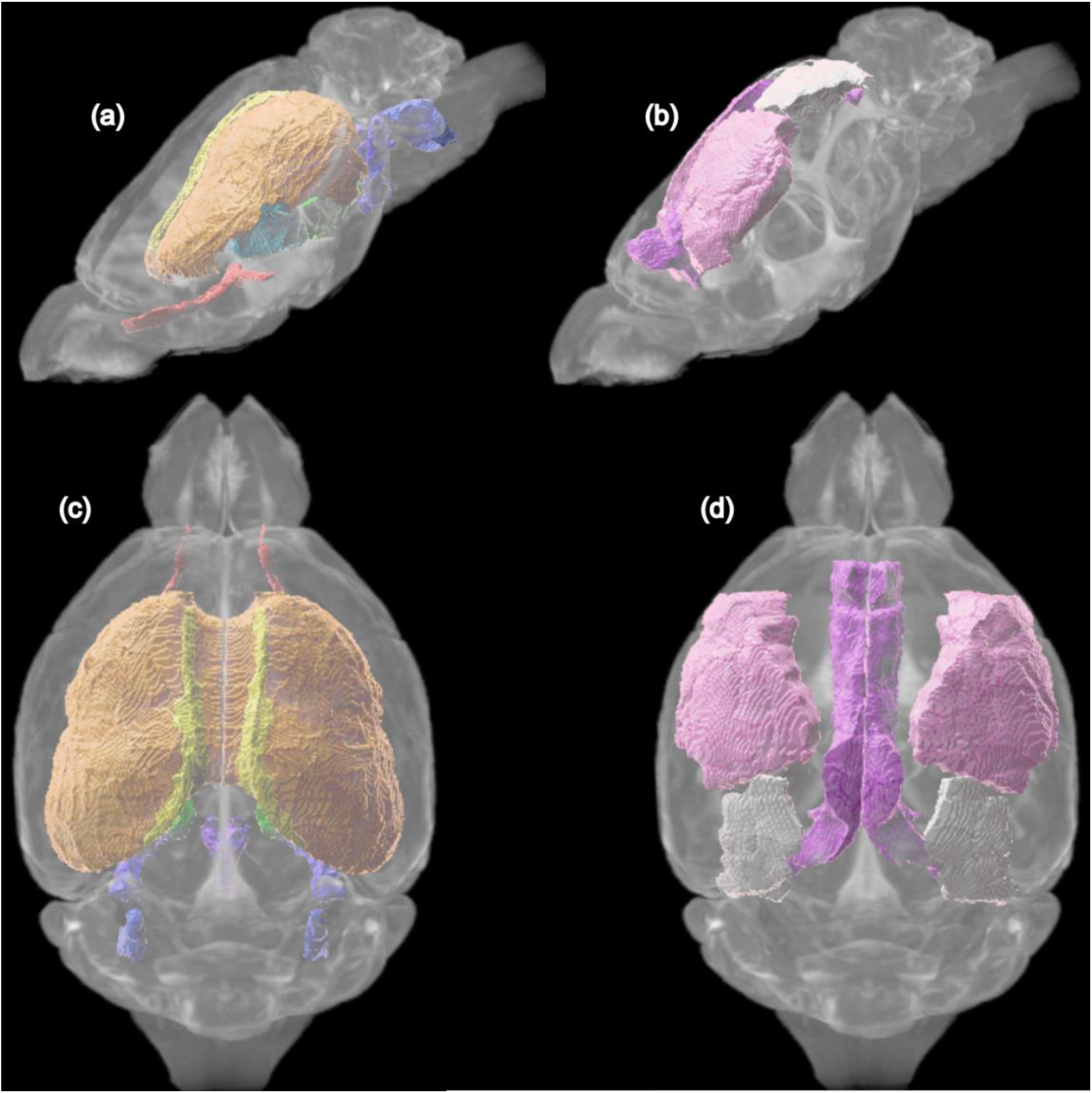
The regions analyzed in this study visualized on an example C57BL/6J specimen with (a and c) showing the white matter regions AC (red), CC (orange), CG (yellow), FI (teal), IC (blue), RN (purple), CP (green), MCP (violet), and ICP (dark blue) viewed from superior and sagittal axis respectively and (b and d) showing the grey matter regions S1 (pink), C (dark pink), V (white) viewed from superior and sagittal axis respectively.

### Characterization of DTI Scalars and Volume Measurements

In order to minimize partial volume effects at the periphery, ImageJ was used to erode each ROI at every time point prior to statistical analysis of the scalar DTI parameter. Figure 3 demonstrates the impact of the erosion on the histogram of FA over three ROI, exemplifying the narrowing and normalization of voxel histograms, and a gradual increase of peak FA with increasing erosion. We chose to use erosion level 1 to balance the removal of edge effects while maintaining an adequate sample of the ROI.

**Figure 3:**
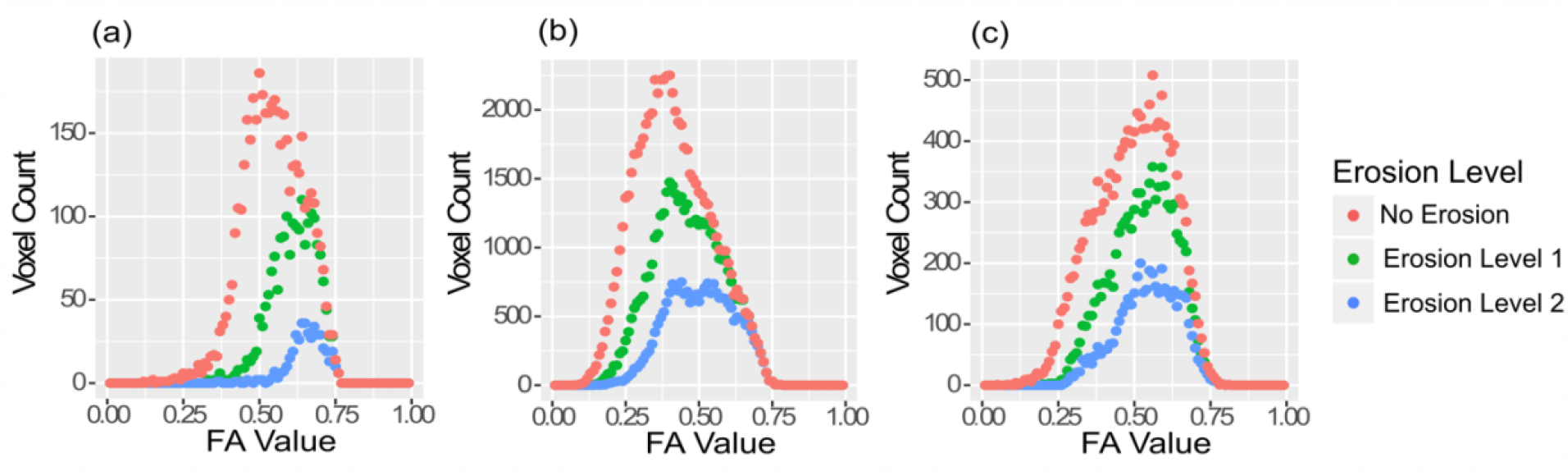
Effects of erosion on the histogram of FA for the (a) anterior commissure, (b) corpus callosum, and (c) internal capsule at the 3 weeks after fixation.

The peak values of each histogram were extracted at each time point as a main study metric. The means and standard deviations of these peaks were the basis of statistical comparison. Data were compared using an ANOVA test to determine if there was any significant variation in time with the given diffusion scalar or volume for the given region-with posthoc pairwise comparison via Tukey’s test to see the most notable time point variations.

The volume variation of each region was calculated in a similar way as the DTI metrics (using an ANOVA test then posthoc pairwise testing). No erosion was applied to the region sizes and the mean percent volume differences, compared to three week time point after fixation, is presented as the key study metric.

### Tractography

DSI studio ^39^ version 20181115 was used to generate tractography for the connectomes. We used the GQI (Generalized Q-Sampling Imaging) model which generates a spin distribution function using a model free reconstruction^40^. Tractography was generated using a tracking plan which terminated after generating 2 million fiber tracts with lengths between 0.5 – 200 mm utilizing all fiber directions at a turning angle of 45 degrees with smoothing and step size of 0.01. The GQI algorithm generates a quantitative anisotropy (QA) analogous to FA. The QA can be normalized by its maximum value of QA, generating the normalized QA (NQA) on a range [0 1]. The NQA threshold for tracking is found for each specimen from rank ordering of the NQA histogram and finding the NQA value corresponding to 10% of the total number of points in the histogram. The mean NQA threshold value for all the specimens over all the times point was 0.105 ± 0.011 (mean ± standard deviation). The connectome values were calculated by counting the number of tracts passing through each region. All connections were included, i.e. there was no thresholding.

Similar methods were used in a recent study of connectome differences between four different strains of mice^9^. In that analysis, approximately five million tracts were generated for each specimen in each strain. In that study, there was little difference in connectomes generated with two million tracts and five million tracts. Thus, we standardized on two million tracts in this new study normalized to an idealized, C57BL/6J brain volume (435 mm^3^) to account for potential changes in brain size across specimens. The mean scale factor is 2.60 +/−0.07 and the average brain volume of this data set is 453 +/−12.0 mm^3^.

The degree of connection - a measurement of the number of connections from one brain region to another - was extracted for all the regions in the atlas ^41^. As noted above, several cortical areas were combined leaving a total number of possible connections at 296. The degree measurement was normalized to this value so that degree of every region ranged between [0 1]. The variation in degree for the 12 example regions is plotted as a function of time after perfusion in the same manner as was used for the scalar DTI metrics.

### Omni-MANOVA

Connectome differences were compared using a graph theory process called OMNI-MANOVA^42^. The connectome is a matrix that expresses the strength of connectivity between each node (i.e. ROI) of the brain. A profile of a specific node, eg. CC, can be constructed based on all the tracts passing through that node. Constructing these profiles allows all the nodes to be expressed as a smaller vector in a common space reducing the complex connectome (332×332 for each specimen) to a more manageable 332x ~3-5 for each specimen based on selecting the principle eigenvalues.

The reduction of terms allows use of multi-variate statistics like MANOVA to determine if the ideally-null tractography fail to reject the null hypothesis between sets of time points. The MANOVA test is a multi-variate form of the ANOVA (a one-way analysis of variance) applied to the node profile described above. We first check that the connectomes are equivalent across the time points and then, by looking at the regions that are significantly different, investigating if that region is known to be problematic, i.e could produce varying tractography due to its structure and features. The p-values corresponding to each region of interest are corrected for multiple comparisons using a Benjamini-Hochberg correction^43^. As we can compare visually the tractography differences, we can utilize less conservative correction methods and then check which nodes are truly different ^44^.This is unlike other statistics applications where the concern for false positive results outweighs benefits of additional contributions obtained by other multiple comparison corrections and where there is no additional check ^45^, such as afforded by our visualization of tractography.

A classical multidimensional scaling was formed in 2-D based on the distances of the Omni-MANOVA embedding tensor via a frobenius norm^46,47^ as a check of how each specimen and group relate to the entire set of study animals. The classical multidimensional scaling is similar to principle component analysis^47^. Rather than focusing on illustrating maximum variation in the set (principle component analysis), the multidimensional scaling illustrates the similarity of the scans. This embedding generates a simplified view, removing region-based analysis of the embedding utilized in the Omni-Manova, so we can make assumptions of the general relationship of our set and compare the spread of groups scanned at different times within the same embedding space.

### Differential Tractography

Once we determined the periods for comparison using the DTI scalar metrics and graph theory based connectomic analysis, we investigated tract differences between the early period (<2 weeks) and the 1 year time point and the stable period using differential tractography in DSI Studio^48^. The method was developed for comparing scans of the same patient at different time points by mapping data sets into a common space and tracking the difference in anisotropy along all the tracks. By looking at the difference in anisotropy, one can determine the stability of the tracking across those two periods. Differential tractography was performed on 4 different specimens in which the 2 week and 8 week time points were compared and the 8 week and 1 year time points were compared.

The initial tracts for differential tractography were generated using DSI Studio version 20200627 to create a series of 50,000 seeds with the same parameters as the initial tracking. We used QA instead NQA, because the percent change criteria in differential tractography is based on QA. The QA thresholding was determined at 10% of the actual QA values. For comparisons between time points, we looked at the changes in QA with four different thresholds, ±20% and ±50%, which had been used previously in a clinical study of demyelination and axonal loss^49^.

## RESULTS

A representative dataset for this work is available on VoxPort (https://civmvoxport.vm.duke.edu/voxbase/studyhome.php?studyid=746). To access the database, a user will need to register an account. Additional data used in this study can be obtained via contact with corresponding author.

### Fractional Anisotropy

The mean peak FA and standard deviation across the specimens at each time point are shown in Figure 4. The mean peak FA of the gray matter regions, Figure 4j-4l with range 0.04-0.24 (−), is substantially lower than the white matter regions, Figure 4a-4i with range 0.23-0.83 (−). The mean peak FA of all ROIs broadly follows an exponential decay, with the most dramatic change from 2-3 days to 2 weeks, decreasing to a plateau from 2 to 8 weeks. For most regions, the FA is relatively unchanged up to a year after fixation.

**Figure 4:**
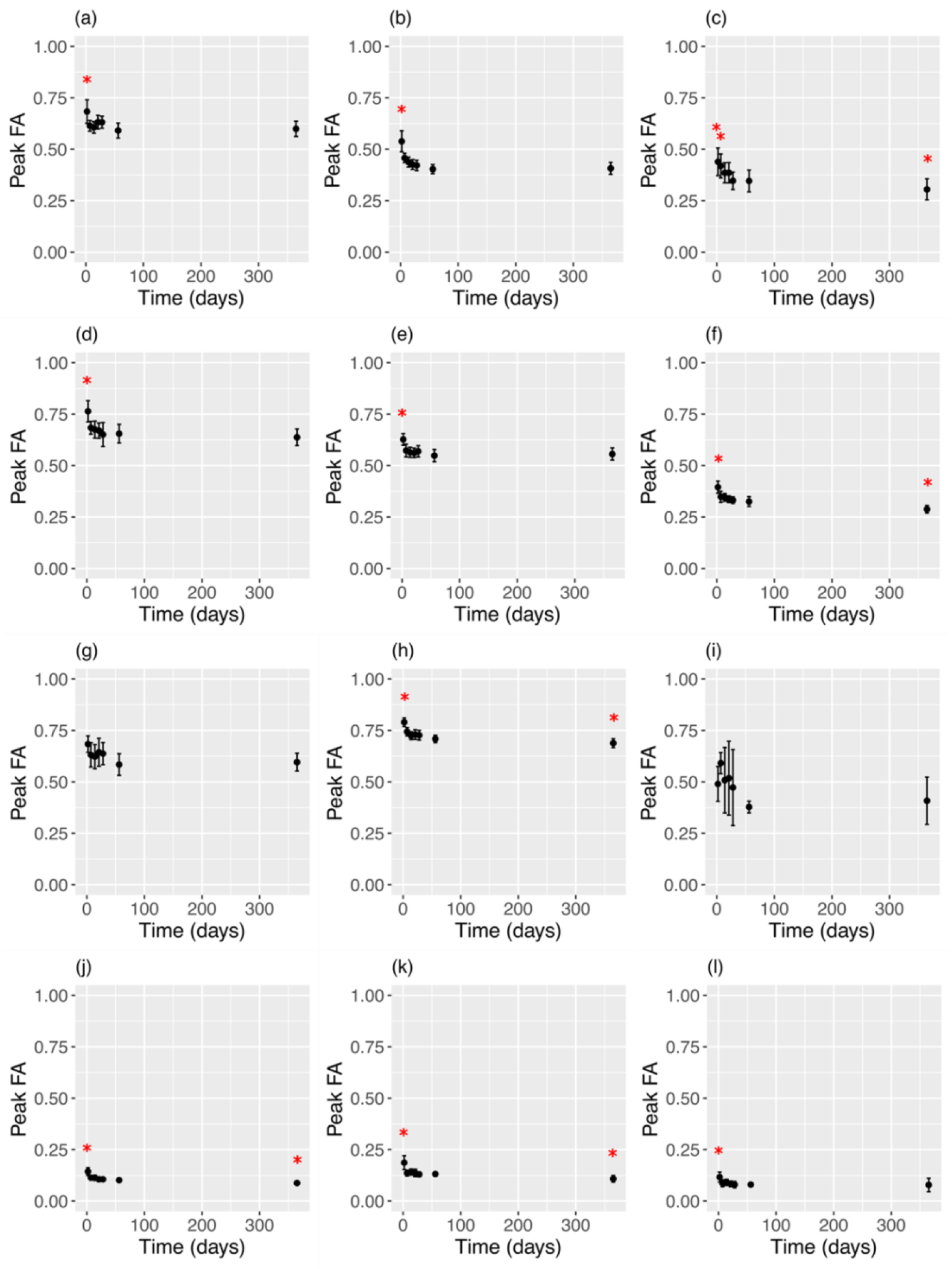
Mean and standard deviation of peak FA across all time points for white matter regions: (a) AC, (b) CC, (c) CG, (d) FI, (e) IC, (f) RN, (g) CP, (h) MCP, and (i) ICP; and gray matter regions: (j) S1, (k) C, and (l) V. Red asterisks denote the time points for a region that differs significantly (p<0.05) in peak FA from at least 2 other time points in the stable period (2 to 8 week post fixation).

An ANOVA test revealed all ROIs changed significantly (p < 0.05) over the entire time period (2-3 days to 1 year). Posthoc Tukey testing (with p < 0.05) determined the intervals of lowest FA variability: the stable time period. All twelve regions showed no significant change in the peak FA from 2 to 8 weeks after fixation.

Ten regions (S1, RN, V, IC, MCP, FI, CC, CG, C, AC) underwent significant changes in peak FA from 2-3 days to 2 weeks after fixation, and nine regions (S1, RN, V, IC, MCP, FI, CC, C, AC) showed a statistically significant difference between at least two points in the stable time period (2 weeks to 1 year) and the 2-3 day period. In Figure 4, significant differences in at least two points in the stable time period are marked by red asterisks for each region. The FA peak of only one region (C) changed significantly from 1 week to 2 weeks and also differed from at least two points in the stable time period. In contrast, the peak values significantly decreased in nine regions (S1, RN, V, IC, MCP, FI, CC, C, AC) from 2-3 days to 1 week after fixation.

The FA in five of the regions (S1, RN, MCP, CG, C) differed between at least two points in the stable period and the one-year time point. But only two of the regions (RN and C) were statistically different from 8 weeks to 1 year. Further investigation revealed that for the five regions that were statistically different at the one year time point, the FA differed by a maximum of 20% relative to the stable period (S1: −18%, RN: −14%, MCP: −5%, CG: −17%, C: −20%), with an overall average change of −8% across all regions.

### Volume

We investigated the volume changes across the 12 selected regions by normalizing to 3 weeks after fixation, which is around the middle of the stable period in the FA. Figure 5 shows the average and standard deviation of the volume change across all time points. There were no statistically discernable changes in volume for any of structures during the stable period with a small increase in the volumes at 1 year after fixation.

**Figure 5:**
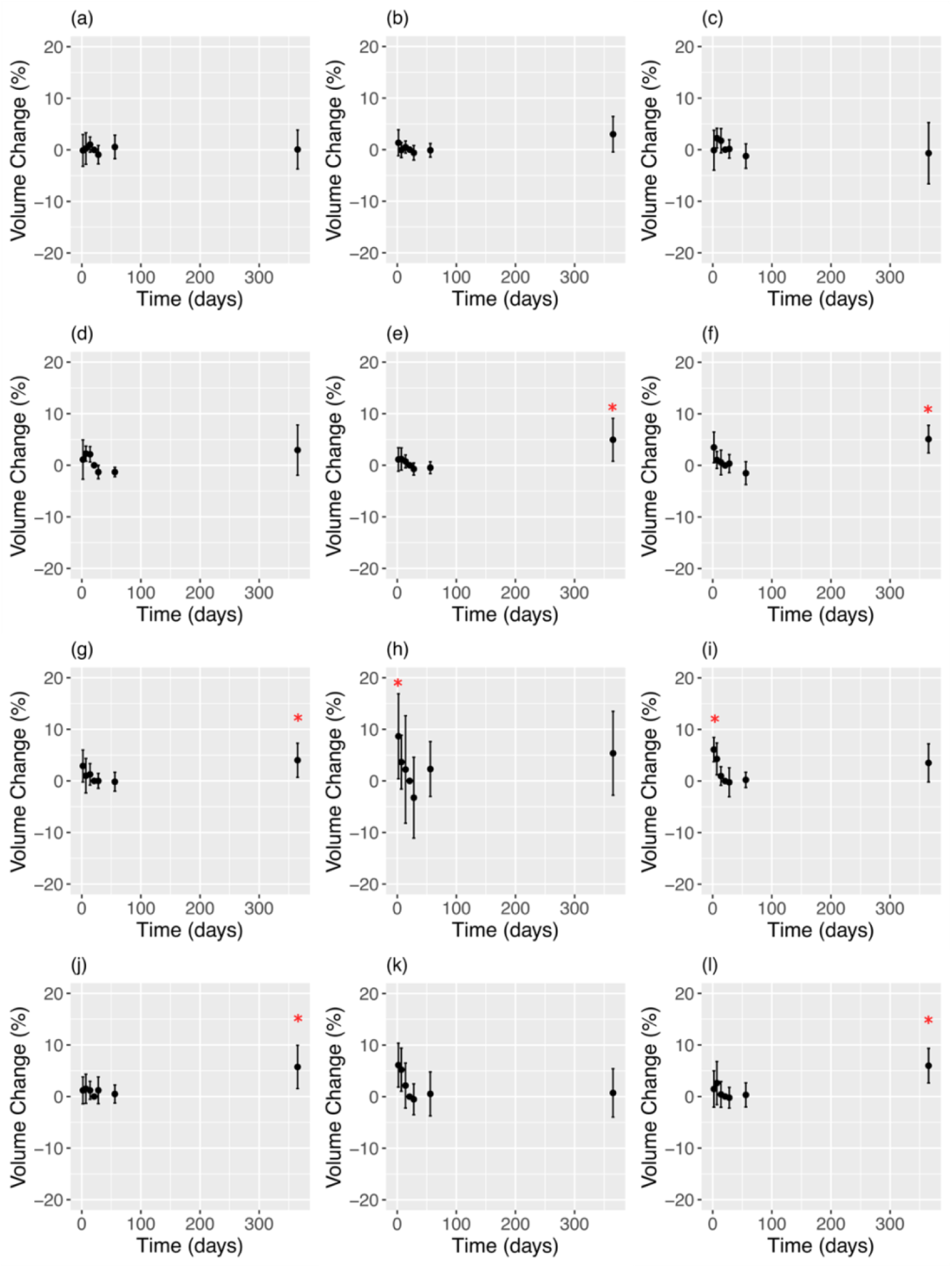
Mean and standard deviations of percent change in volume compared to the 3-week time point across all scans for white matter regions: (a) AC, (b) CC, (c) CG, (d) FI, (e) IC, (f) RN, (g) CP, (h) MCP, (i) ICP; and gray matter regions: (j) S1, (k) C, (l) V. Red asterisks denote the time points at a region that significantly (p<0.05) differs in volume (mm^3^) from at least 2 other time points in the stable period (2 to 8 weeks post fixation).

An ANOVA test determined that eight of the twelve regions had statistically significant change in volume across time (p < 0.05; S1, RN, V, IC, MCP, ICP, C, CP). The four regions without significant volume differences over time (FI, CC, CG, AC) were white matter regions. Post-hoc testing was performed on the eight regions that changed significantly (p < 0.05) in volume over time in order to discern the time period in which the volume is stable. Similar to the results for the diffusion metrics, there were no significant changes in volume between 2 and 8 weeks after perfusion. Outside of this stable period, we see with the pairwise comparison that volumes at the one year time point varied significantly from the stable period in five of the eight regions (S1, RN, V, IC, CP). While the number of regions with volumes that were different at one year is large, the percent difference of volume is small, with a mean change of 5% (S1: 5%, RN: 5%, V: 6%, IC: 5%, CP: 4%). Unlike the FA, only the volumes of two of the eight regions (MCP, ICP) differed significantly at the 2-3 day time point compared to at least 2 other time points in the stable period. No volume of any region at 1 week after fixation differed significantly from at least two time points in the stable period nor significantly differed from the volume at the 2 week time point. In Figure 4, the significant changes from at least two points in the stable time period are marked by red asterisks for each region.

### Supplemental Scalar Metric Analysis

Four additional scalar metrics (mean diffusivity (MD), radial diffusivity (RD), axial diffusivity (AD), and normalized degree) were analyzed for this work in the same way as FA and volume. The resulting figures and analysis are included within the supplement. The results for those additional metrics follow that of the FA and volume.

### Omni-MANOVA

The Omni-MANOVA analysis in Figure 6 provides an expanded analysis of the stability of the connectome by showing the reduced vector comparisons at multiple time points. In Figure 6a, in which all the time points are compared, there is a marked separation of the 2-3 day and 1 week data (red and green) from the rest of the groups (blue, cyan, magenta, yellow and black). This indicates the 2, 3, 4, 8 week, and 1 year measurements are more similar to one another than the early time points. The rank ordered p-value results of the MANOVA test, identified 160 of the 332 regions that reject the null hypothesis that the connectome is unchanged, i.e. the vectors describing connections of 160 nodes may have changed. In Figure 6b, the 2-3 day and 1 week time points have been removed. The 1 year time point does not overlap the stable period as closely as in Figure 6a but the difference is less than one standard deviation of the constituent groups and the reconstruction covers less area showing that the data are more similar than all the time groups together. The MANOVA test indicates that profiles (vectors) of 4 out of the 332 regions of interest reject the null hypothesis, Table 1. All are cortical and none are bilateral. Boundaries of these regions are poorly defined in our atlas so the use of FA as an automated mapping criterion results in higher variability in mapping. Suspecting this variation is due to mainly the boundaries of the included data within the embedding, 2 weeks and 1 year, we continue by trying two different embeddings each removing one of these boundary time points. In Figure 6c, the 2-week data has been removed, the plot shows 3 weeks to 1 year. There is little change from Figure 6b and no determined significant changes in connectivity. This helps us conclude that the stable period is 2-8 weeks. Figure 6d plots the data for this 2-8 week stable period. Note the spread (standard deviation) in this plot is small and not that different than Figure 6c. In addition, there were no profiles which had changed significantly. This suggests that the 1 year data is not drastically different from that gathered during this stable period.

**Table 1:** The regions associated with 2, 3, 4, 8 week and 1 year embedding that failed to reject the null hypothesis according to BH correction, sorted by raw p-value.

| Region Abbreviation | Region Name | Hemisphere |
| --- | --- | --- |
| V1B | primary visual cortex, Binocular Area | Right |
| DLEnt | dorsolateral entorhinal cortex | Right |
| A24aPrime | cingulate cortex, area 24a prime | Right |
| S1FL | primary somatosensory cortex, forelimb region | Left |

**Figure 6:**
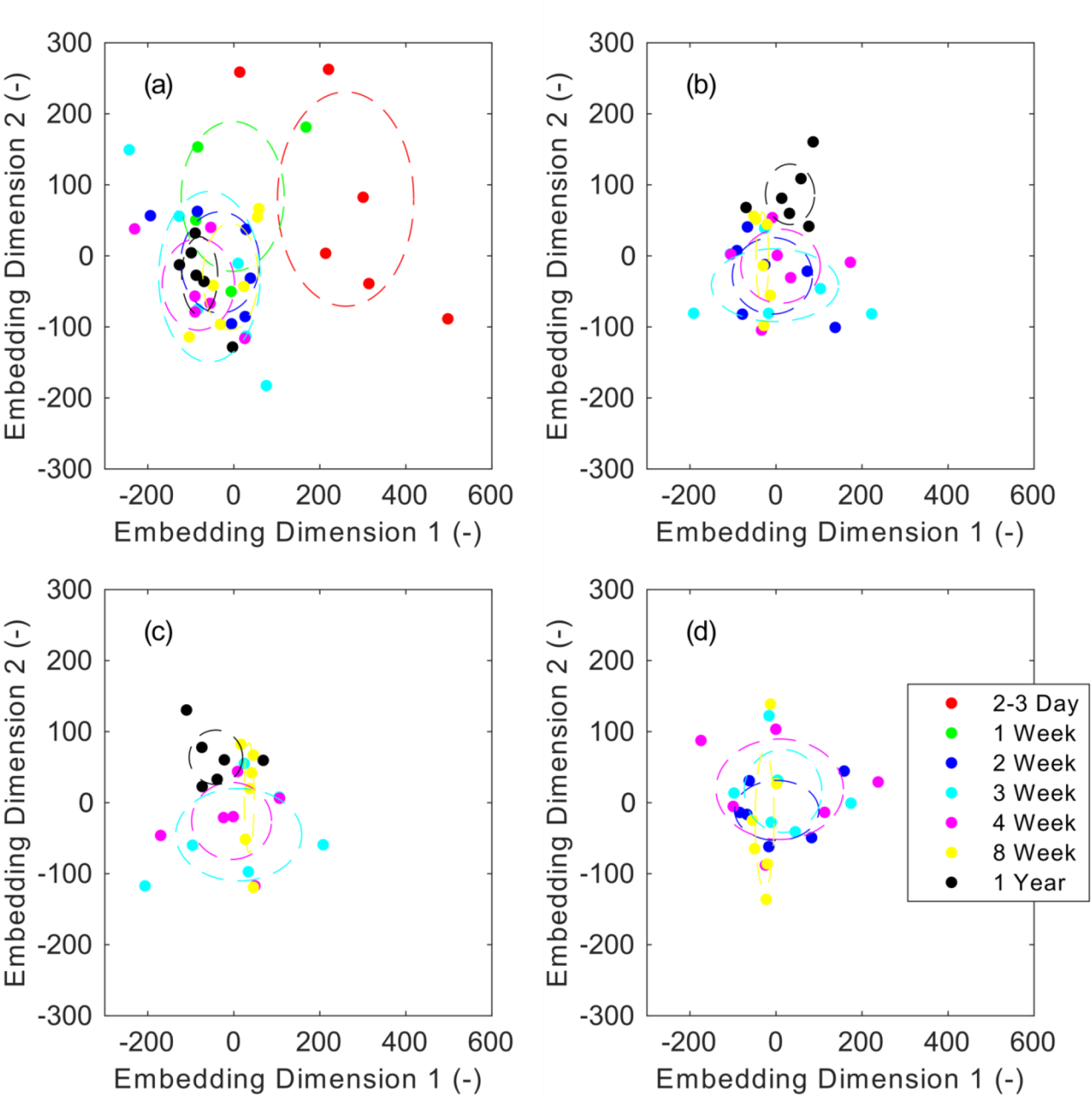
Reconstructions of the adjacency spectral embedding created using Omni-MANOVA framework resulting from (a) all time points, (b) 2, 3, 4, 8 week, and 1 year time points, (c) 3, 4, 8 week, and 1 year time points, and (d) 2, 3, 4, and 8 week time points. The dotted lines are centered on the mean of individual points in the specific time periods with the spread corresponding to the standard deviation in embedding dimension 1 and 2. Individual points in the plots represent each specimen in this study.

### Differential Tractography

Differential tractography was employed as a final test of tissue stability. We compared four specimens at 2, 8 week, and 1 year time points. The percent change in QA at 2 weeks and 1 year utilized the 8 week scans as the base point. Using 50,000 seeds with whole brain tracking, the average track number and length for the groups were nominally the same, 21482 +/−240 and 5.03 +/−0.111 mm respectively, at the three time points, indicating that there is not a drastic change in the diffusivity on which the tractography is based. The average numbers associated with each time point are shown in Supplemental Table 1.

When QA *change* is used as the thresholding condition, the mean tract length is approximately a fifth of the tract length in the QA images. The mean tract number and mean tract length at ±20% and ±50% thresholds are shown in Supplemental Table 2. The differential tracts are small, scattered, and likely are related to edge effects. Differences between the 2 and 8 week scans are negligible. The differences between the 8 week and 1 year scan are most notable at −20% suggesting some systematic though small reduction in FA as the tissue rests in PBS for a year. One can relate the differential tracts to the global length and density of tracts comparison to all tracts passing through the left portion of the CC for the same specimen (180423-2:1) at the three time points from the 50,000 seeded whole brain track data. We picked 180423-2:1 because in the whole brain tracking, the response of number of tracts and track length was most close to the average result. Example images of this tractography at all three time points are shown in Figure 7b-7d. Both Figure 7b and Figure 7d used Q-Space Diffeomorphic Reconstruction^50^ (QSDR) to place the 2 week and 1 year data into the same orientation alignment as the 8 week data (Figure 7c), and thus needed to use the 8 week label for segmentation of CC tracts. We see more grouping of key tracts, such as the arching structure of the CC and projections into the olfactory bulb, than illustrated by the difference tractography. As a comparison, the average number of tracks shown in Figure 7b-7d is 2520 +/−60 which is approximately the same number of average tracks found for all specimen at −20% threshold comparison of 1 year and 8 weeks. Interestingly, while the number of tracts was the same, the average tract length of Figure 7b-7d is 11.5 +/−0.796 mm which is an order of magnitude longer than the multi-specimen averaged −20% threshold comparison of 1 year and 8 weeks. Thus, this confirms that our remaining differences in tractography shown in the differential tractography are negligible on the whole.

**Figure 7:**
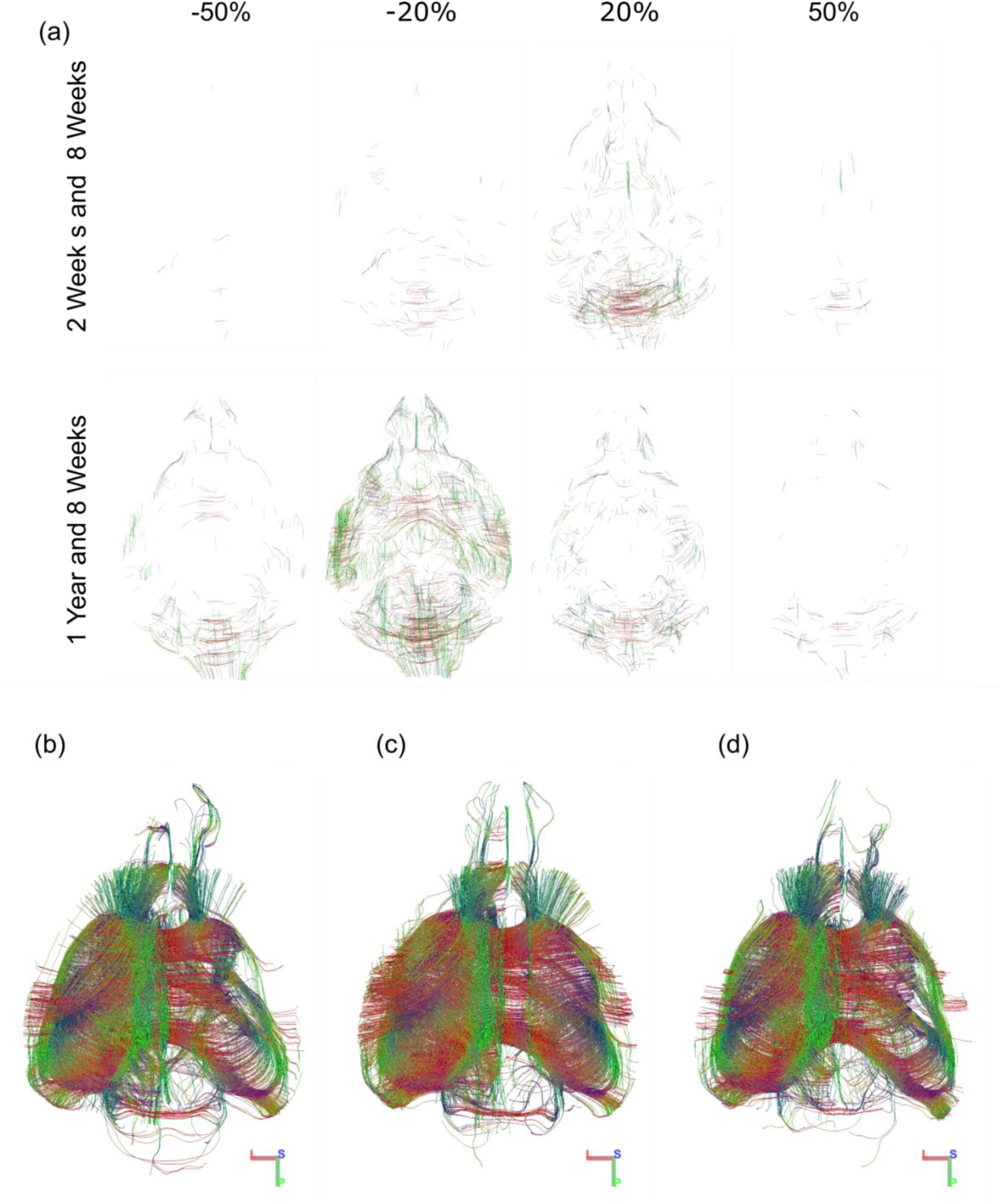
A comparison of (a) differential tractography measuring the change of QA at 2 weeks and 1 year using 8 weeks as a basis and tracking parsed to only those that pass through the CC at (b) 2 weeks (c) 8 week, and (d) 1 year post fixation. All images are produced with specimen 180423-2:1 which had the largest difference in track length and number at ±20% and ±50% in the differential QA data.

## DISCUSSION

Through our analysis of fractional anisotropy, axial diffusivity, radial diffusivity, and mean diffusivity, we have sought to understand the changes in diffusion tensor scalars due to fixation over time. Similar to the literature results for T1 and T2 changes post-fixation ^21 22,51 20^, we found that some of the DTI scalar values varied significantly during the first two weeks following perfusion fixation, but eventually stabilized to a nominally constant value. Changes in the diffusion metrics were statistically insignificant between 2 and 8 weeks providing a stable period. Specimens scanned during this window can be compared without confound from fixation. Conversely the scalar DTI metrics in multiple gray and white matter regions changed significantly over the first two weeks of fixation suggesting our standard operating procedure should include a stabilization period of at least 2 weeks prior to scanning.

There are statistically detectable differences in the scalar metrics between the stable period and 1 year. But the DTI parameters at one-year post fixation have changed only marginally from the measurements during the stable period. The largest change was in the AD at 13% with an average change of 7% in AD across all the structures measured. This suggests that specimens may be reexamined for a longer period without serious confound to the study.

Volume of the 12 structures was statistically unchanged during the 2 to 8 week stable time period. Volume change out to one year post fixation was ~ 3% over all the regions. De Guzman et al have performed a nicely detailed study of volume changes using protocols similar to ours (i.e. perfusion fixation, 1-5 day formalin immersion, long term storage in saline or water)^52^. The period of observation (to 150 days) is comparable to ours. They found a rapid change in volume (3.5%/day) when tissue is stored in formalin and ~3%/month when tissue is stored in PBS. Our data is consistent with theirs though the time points of comparison are not identical. De Guzman showed volume changes at 5 months of white matter ranging between ±4% and grey matter between −4% and +5%. For our data, the maximum volume change in any volume at 1 year is 6% and an average change of 3% over all regions. While small, these changes are still larger than the whole brain volume change, which increases to a maximum of 2% volume change from 2 weeks to one year. But it bears repeating that during the stable period (2-8 weeks) there were no statistically discernable changes in volume in the 12 structures surveyed. In a recent publication, we measured the coefficient of variation between left and right hemispheric structures in four different strains of mice as a measure of technical error ^9^. That coefficient of variation was <5% for structures with volumes > 1 mm^3^. This variation is probably an indication of the reproducibility of the registration pipeline.

The Omni-MANOVA results further are consistent with changes in the scalar metrics during the first two weeks and stable time period. The points associated with the early (2/3 day and 1 week) time period stand out as particularly different than the other groups. Figure 6a maps all the data into a common space. In this mapping, there are 160 of 322 regions with connectivity that differs. In Figure 6b, the 2/3 day and 1 week data have been removed and there are only 4 regions (all cortical) that differ with p values that are significant. The connectomes are remarkably stable from 2 weeks out to one year. Similar to the conclusion derived from the parametric analysis of the scalar DTI metrics, the one year connectome data is similar enough to the stable period that the one year time point can be considered in practice invariant from the stable time period.

The differential tractography results also confirms the stability of the tissue over the 2-8 week period with minor changes between 8 weeks and 1 year. There is a suggestion of increased anisotropy (+20%) particularly in the cerebellum in comparison to the 2 and 8 week data but the mean track length and number (0.91 mm, 570 tracks) in the difference image vs mean track length and number (5 mm, 21516 racks) in the base images suggest this is probably an edge effect. The mean track length in the 8 week and 1 year difference image is also small (0.97mm @ +20%) suggesting an edge effect. But there is an increased in number and mean length at −20% from 0.82 mm with 111 tracks @ −20% to 1.18 mm with 2490 tracks when comparing 8 weeks and 1 year. This is consistent with the bulk scalar metrics in which the AD at one year was statistically different from the stable period in eight of the twelve regions (Figure S3).

## CONCLUSION

Numerous researchers have studied the effects of autolysis, fixation (post mortem interval after death, length and type of fixative and storage (in fixative, water, saline) on the diffusion properties of tissue ^19,24,25,27,30,53–55^. Our *very specific protocol for quantitative connectomics in the rodent brain* has been constructed with the results of these studies in mind. For example, it is clear that fixation is essential to limit autolysis. Perfusion fixation is considerably more effective than immersion. Extended fixation (>24 hrs) causes continuing reduction in T2 (and therefore reduced signal to noise). Storage in buffered saline is desirable to minimize tissue swelling/changing. And active staining ^16,56^, i.e. perfusion with a contrast agent is essential for microscopic imaging. Finally, our application differs considerably from much of the previous literature. We are scanning the mouse brain. Perfusion fixation is easily performed which is generally not possible in the human studies. We are doing so with spatial resolution (45 um^3^ i.e 91 pl). The spatial resolution of DTI in clinical MRI is generally >1 mm^3^, i.e. a difference of nearly 11,000 × in voxel volume.

Quantitative connectomics is now being applied in both clinical and basic science settings. It provides remarkable insight into brain structure and health. But is has limitations. Acquisition protocols differ widely in spatial resolution, angular sampling, gradient weighting, and processing algorithms. Maier-Hein and a consortium of many of the leaders in the clinical use of connectomes raised significant concerns over false positive connections even with optimized clinical protocols and pipelines ^57^. And as noted above, comparison of clinical protocols with methods for the mouse brain is fraught. Retroviral tracers continue to be the gold standard for quantitative differentiation of afferent and efferent connections in the mouse brain ^58^.Yet diffusion derived connectomes at microscopic resolution provide intriguing applications that cannot be done with tracers ^9,13,59^. But like any new methodology, the protocols must be optimized and the limits understood. We have provided a carefully constructed comparison of our protocol with retroviral tracers ^60^. We have optimized that protocol for routine use ^34^. And in this work, we have begun to define how to use it with confidence.

## Supporting information

Graphical Abstract

## ACKNOWLEDGEMENTS

This work was supported by the NIH/NIBIB National Biomedical Technology Resource Center P41 EB015897 (to GA Johnson), NIH 1S10OD010683-01 (to GA Johnson) and NIH 1R01NS096720-01A1 (to GA Johnson). We are grateful to Lucy Upchurch for technical support and Tatiana Johnson for assistance in preparing the manuscript.

## ABBREVIATIONS AND UNITS

(ROI): Selected Regions of Interest
AC: anterior commissure
CC: corpus callosum
CG: cingulum
CP: cerebral peduncle
FI: fimbria
IC: internal capsule
ICP: inferior cerebral peduncle
MCP: middle cerebral peduncle
RN: raphe nucleus
C: cingulate cortex
S1: primary somatosensory cortex
V: primary visual cortex
DTI: (Diffusion Tensor Imaging) Measure Abbreviations:
FA: fractional anisotropy [unitless]
AD: axial diffusivity [mm^3^/s]
RD: radial diffusivity [mm^3^/s]
MD: mean diffusivity [mm^3^/s]

[Dataset] Jaclyn Xiao, Kathryn J. Hornburg, Gary Cofer, James J. Cook, Yi Qi, and G Allan Johnson; 2020; “Stained Time-course Study Representative Data”; VoxPort; https://civmvoxport.vm.duke.edu/voxbase/studyhome.php?studyid=746

