## Supplementary material for "A time-course study of actively stained mouse brains: DTI parameter and connectomic stability over one year": Graphical Abstract

This test/retest diffusion weighted (DW) MRI connectomics study determines the time period post-fixation that an actively perfused mouse brain (C57BL/6J) can be remeasured and still maintain consistency in result. We determined from a mix of analysis of connectomics and DW metrics that there are three distinct time periods: early (prior to 2 weeks), stable (2 weeks to 8 weeks), and late (post 8 weeks); with the early time period being the most unstable of the three time periods.

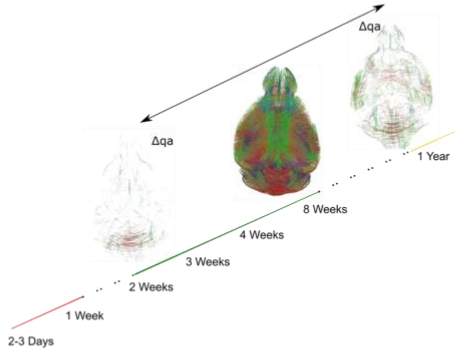
