## Supplemental Data for "A time-course study of actively stained mouse brains: DTI parameter and connectomic stability over one year"

**SUPPLEMENT**
Supplemental Scalar Metric Analysis

In addition to fractional anisotropy (FA) and volume, four additional scalar metrics were investigated within this work: mean diffusivity (MD), radial diffusivity (RD), axial diffusivity (AD), and normalized degree. Degree is a scalar metric relating to how many other regions the ROI is connected to, much like the DTI metrics which we also discuss here. The graphs of the average peak value and standard deviation for each time point are included within this supplement (MD—Supplemental Figure 1, RD – Supplemental Figure 2, AD—Supplemental Figure 3, and normalized degree – Supplemental Figure 4). Generally, there is most instability at the early (<2 week) time point with a stable (flat) period from 2 to at least 8 weeks.

Using ANOVA testing, we determined if the response across the 12 regions varied with time ($p<0.05$) for each metric. For MD and RD, all 12 regions significantly varied with time. For AD, 10 regions significantly varied across time. The unchanged regions associated with AD are ICP and FI. Only 7 of the 12 regions (RN, IC, FI, CC, CG, CP, AC) significantly differed for the normalized degree metric.

Using a posthoc Tukey's test on the regions identified by the ANOVA test as significantly varying with time, we discovered which time combinations provided significant instability ($p < 0.05$) for each scalar metric. In all the regions identified in the ANOVA, the 2 to 8 week time period was rarely determined to have changed significantly and thus is the "stable time zone". The only significant change combinations for pairs between 2 to 8 weeks are MD: C [2 versus 8]; RD: FI [2 versus 8 and 3 versus 8], S1 [2 versus 8], V [2 versus 8], ICP [2 versus 8]. Those regions are mainly cortical which have the most possibility of variable boundaries, as previously discussed. A stability metric has been defined as significant changes from at least two points in the "stable time zone". The time points which fail this criteria are marked by red asterisks for each region in the Supplemental Figures 1-4.

Investigating the earlier time point comparisons with the posthoc results, there is much more instability than the 2 to 8 week period, but a trend towards increased stability at the 1 week timepoint. For MD, all regions at 2-3 days and 5 regions (RN, V, ICP, C, AC) at 1 week differed from at least two timepoints in the stable period. For RD, all regions at 2-3 days were deemed changed from stable time points, while at 1 week only 5 of the 12 (S1, RN, V, MCP, and F) significantly varied based on our stability condition. For AD, 7 out of 10 regions (S1, RN, V, MCP, CC, CG, C) at 2-3 days after fixation and one region (V) at the 1 week timepoint differing from at least 2 timepoints in the stable period. For normalized degree, the same pattern of all regions significantly varying based on the stability condition at 2-3 days continued with no regions determined under the same criteria at 1 week.

Investigating the later timepoint, we note some instability, but that instability tends to be small. For MD, 7 regions (RN, IC, MCP, ICP, CC, CP, AC) varied at 1 year based on the stability criteria. Although, the percent change of peak MD (1 year compared to average stable time period value) is small, equating to a maximum value of 10% (AC and CC). For RD, 4 regions (V, IC, MCP, ICP) are deemed unstable. Again, the changes in peak RD values at one year compared to the stable period are small, with a maximum percent change of 6% (IC, MCP). The AD values were more variable than FA, MD, and AD at the one year time point with 8 of the 10 regions differing based on the stability criteria and a maximum change at the 1 year point to the stable period of 13% (MCP). Normalized degree is the most stable at 1 year with only 1 region (RN) with significant differences from 2 other stable region time points.

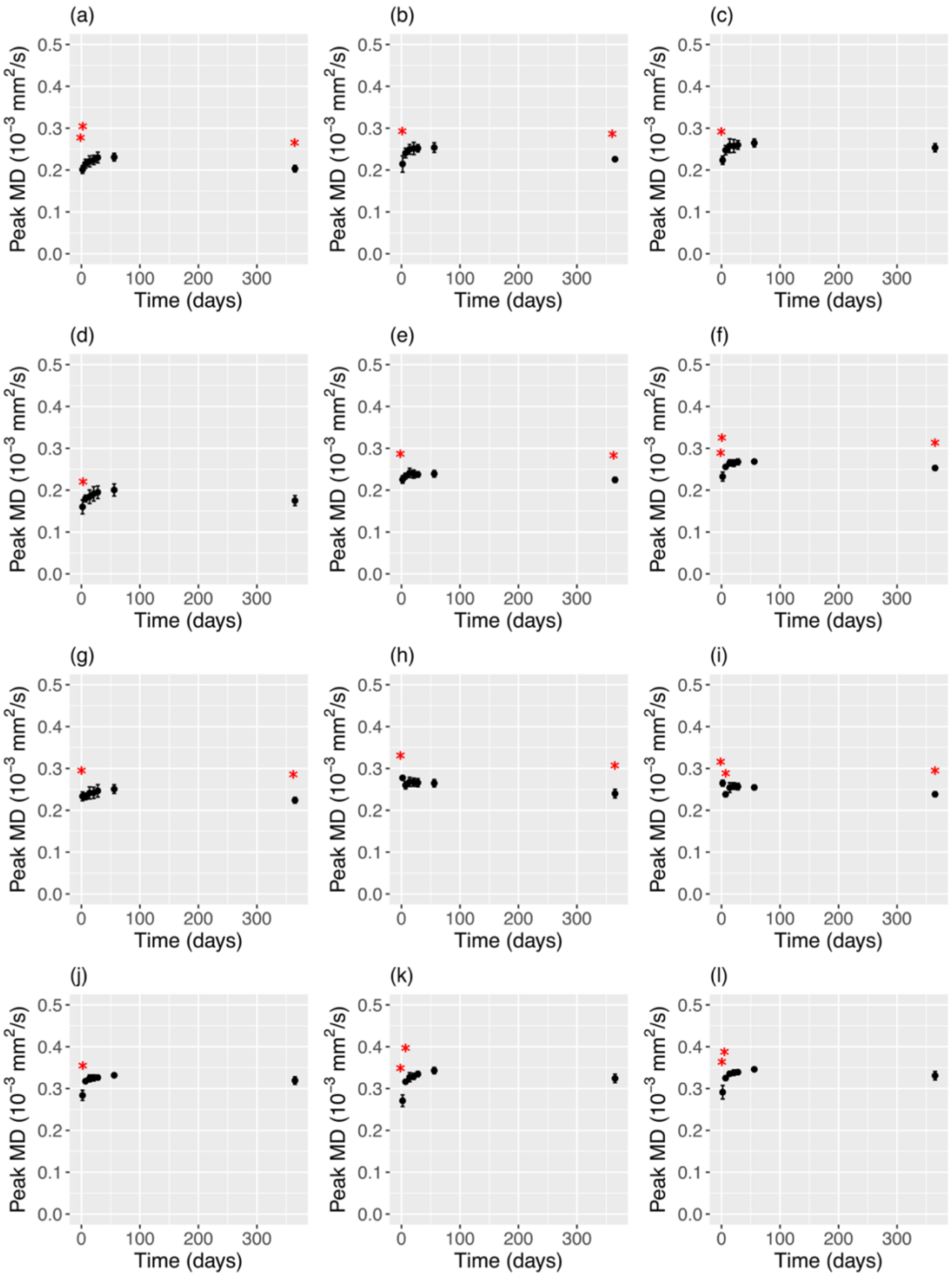


Supplemental Figure 1: Mean and standard deviations of MD peak values across all specimen at all time points (2-3 days, 1 week, 2 weeks, 3 weeks,4 weeks, 8 weeks, and 1 year after fixation) for white matter regions: (a) AC, (b) CC, (c) CG, (d) FI, (e) IC, (f) RN, (g) CP, (h) MCP, (i) ICP; and gray matter regions: (j) S1, (k) C, (l) V. Red asterisks denote the timepoints at a region that significantly ($p<0.05$) differs in peak MD value from at least 2 other timepoints in the visually stable period (2 to 8 weeks post fixation).

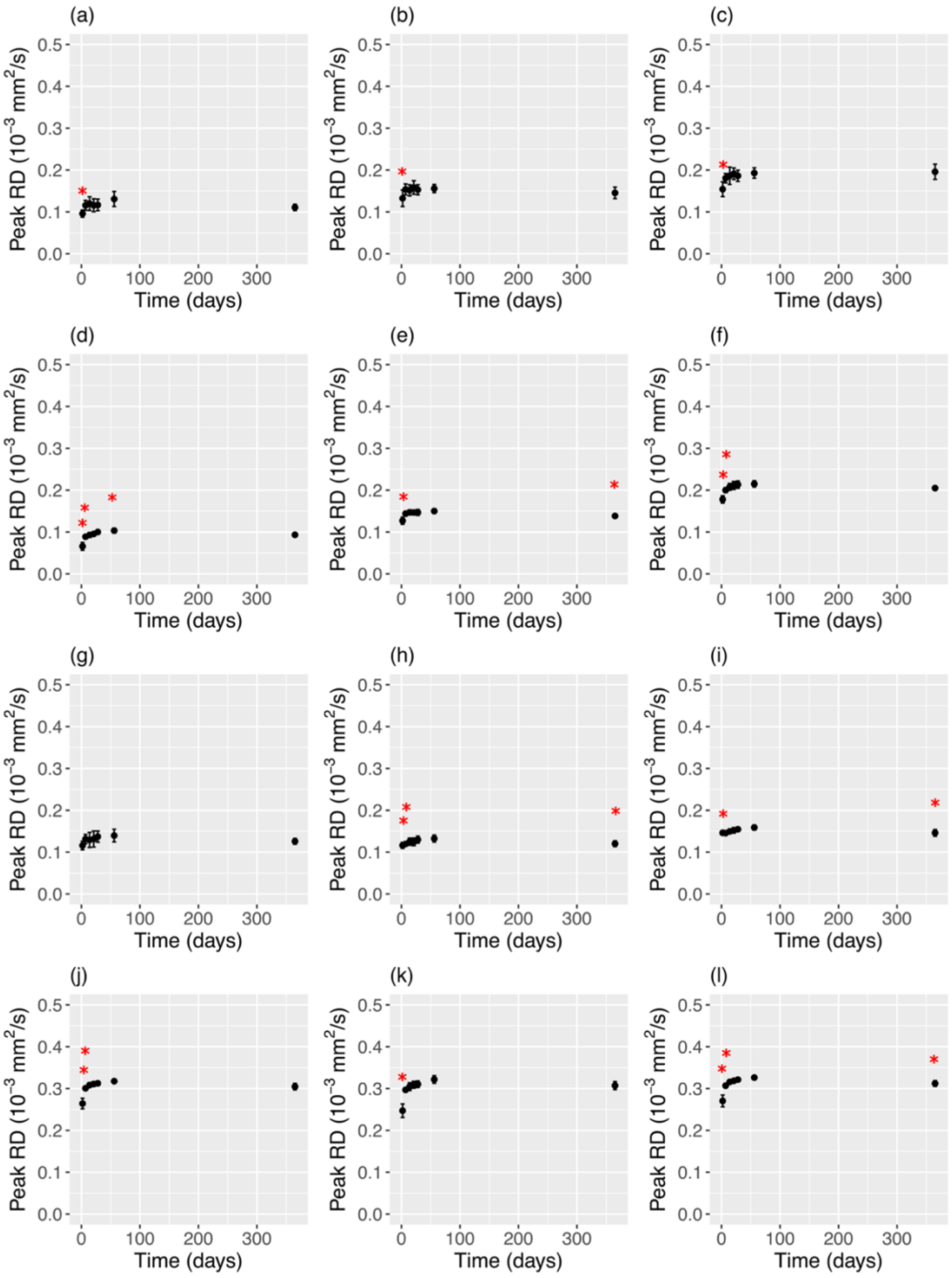


Supplemental Figure 2: Mean and standard deviations of RD peak values across all specimen at all time points (2-3 days, 1 week, 2 weeks, 3 weeks, 4 weeks, 8 weeks, and 1 year after fixation) for white matter regions: (a) AC, (b) CC, (c) CG, (d) FI, (e) IC, (f) RN, (g) CP, (h) MCP, (i) ICP; and gray matter regions: (j) S1, (k) C, (l) V. Red asterisks denote the timepoints at a region that significantly ($p<0.05$) differs in peak RD value from at least 2 other timepoints in the visually stable period (2 to 8 weeks post fixation).

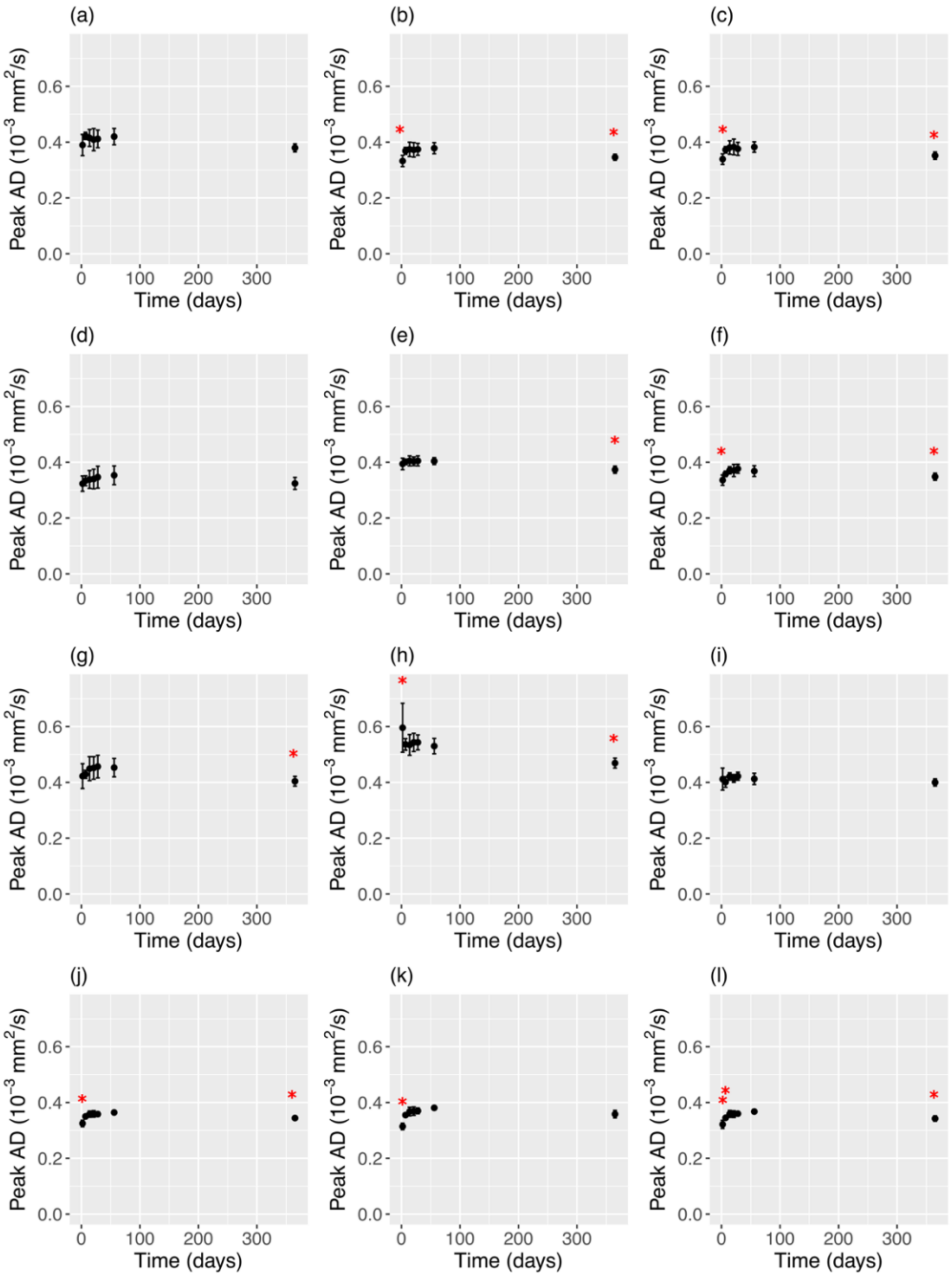


Supplemental Figure 3: Mean and standard deviations of AD peak values across all specimen at all time points (2-3 days, 1 week, 2 weeks, 3 weeks, 4 weeks, 8 weeks, and 1 year after fixation) for white matter regions: (a) AC, (b) CC, (c) CG, (d) FI, (e) IC, (f) RN, (g) CP, (h) MCP, (i) ICP; and gray matter regions: (j) S1, (k) C, (l) V. Red asterisks denote the timepoints at a region that significantly ($p<0.05$) differs in peak AD value from at least 2 other timepoints in the visually stable period (2 to 8 weeks post fixation).

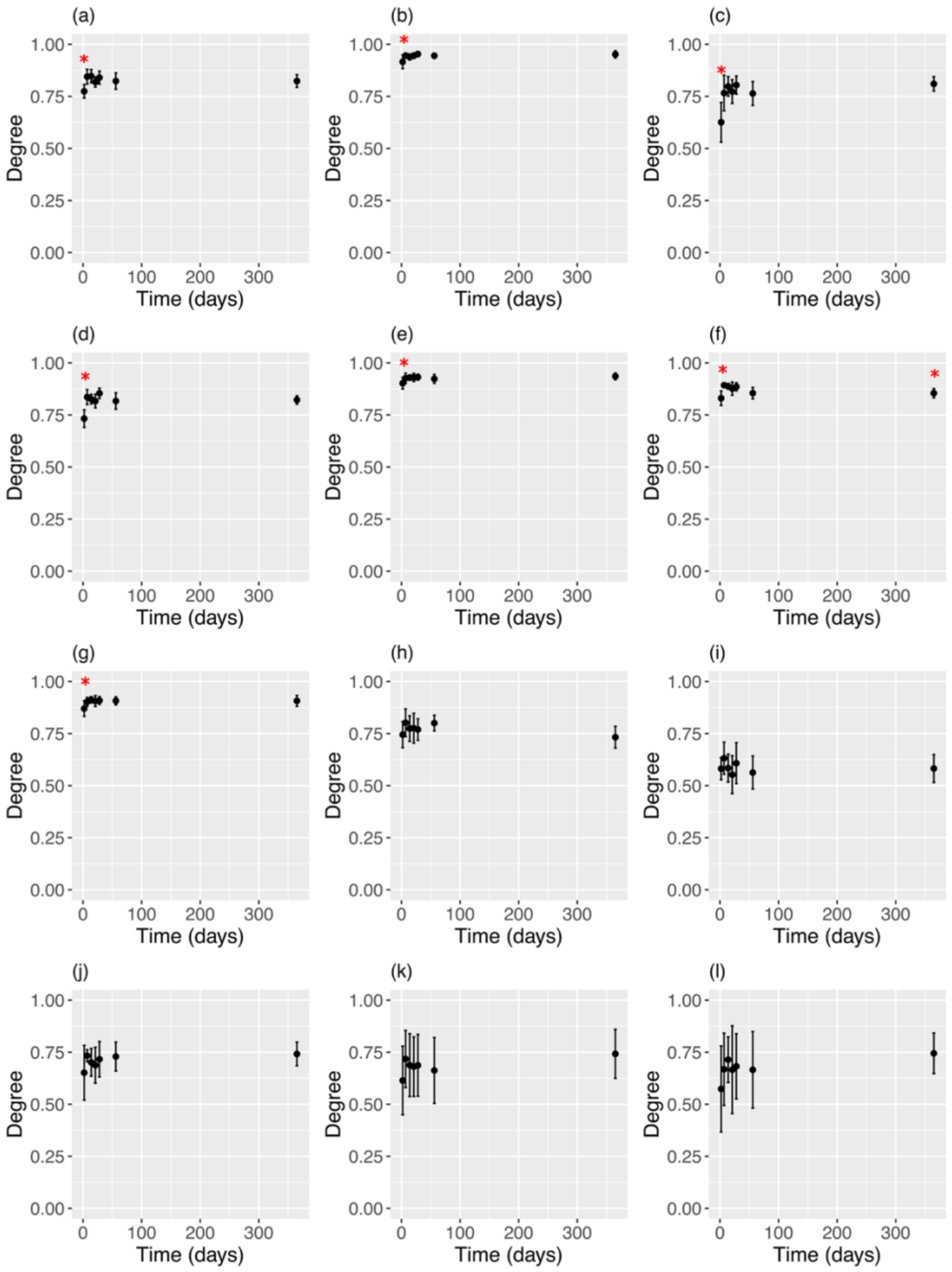


Supplemental Figure 4: Mean and standard deviations of normalized degree values across all specimen at all time points (2-3 days, 1 week, 2 weeks, 3 weeks, 4 weeks, 8 weeks, and 1 year after fixation) for white matter regions: (a) AC, (b) CC, (c) CG, (d) FI, (e) IC, (f) RN, (g) CP, (h) MCP, (i) ICP; and gray matter regions: (j) S1, (k) C, (l) V. Red asterisks denote the timepoints at a region that significantly ($p<0.05$) differs in normalized degree values from at least 2 other timepoints in the visually stable period (2 to 8 weeks post fixation).

Supplemental Table 1: Comparison of Average Mean Tract Number (rounded to nearest whole tract) and Mean Tract Length for the Two Week, Eight Week, and 1 year Time Groups in a 50,000 Tract Seeding Experiment

| Time Group | Mean Tract Number | Mean Tract Length (in mm) |
|---|---|---|
| 2 Week | 21604 | 5.09 |
| 8 Week | 21516 | 4.98 |
| 1 Year | 21328 | 5.04 |

Supplemental Table 2: Comparison of Average Mean Tract Number (rounded to nearest whole tract) and Mean Tract Length with Eight week as Basis Time Period in a 50,000 Tract Seeding Experiment of Percent Change of QA

| Time Group | Change in QA (Mean Tract Number, Mean Tract Length) | | | |
|---|---|---|---|---|
| | -50% | -20% | 20% | 50% |
| 2 Week and 8 Week | (13, 0.759 mm) | (111, 0.821 mm) | (570, 0.907 mm) | (54, 0.803 mm) |
| 8 Week and 1 Year | (468, 0.985 mm) | (2490, 1.18 mm) | (701, 0.967 mm) | (108, 0.803 mm) |
